# SARS-CoV-2 Nsp1 binds ribosomal mRNA channel to inhibit translation

**DOI:** 10.1101/2020.07.07.191676

**Authors:** Katharina Schubert, Evangelos D. Karousis, Ahmad Jomaa, Alain Scaiola, Blanca Echeverria, Lukas-Adrian Gurzeler, Marc Leibundgut, Volker Thiel, Oliver Mühlemann, Nenad Ban

**Affiliations:** Department of Biology, Institute of Molecular Biology and Biophysics, ETH Zurich, Zurich, Switzerland; Department of Chemistry and Biochemistry, University of Bern, Bern, Switzerland; Institute of Virology and Immunology, Bern, Switzerland; Department of Infectious Diseases and Pathobiology, Vetsuisse Faculty, University of Bern, Bern, Switzerland

## Abstract

The non-structural protein 1 (Nsp1), also referred to as the host shutoff factor, is the first viral protein that is synthesized in SARS-CoV-2 infected human cells to suppress host innate immune functions^1,2^. By combining cryo-electron microscopy and biochemical experiments, we show that SARS-CoV-2 Nsp1 binds to the human 40S subunit in ribosomal complexes including the 43S pre-initiation complex. The protein inserts its C-terminal domain at the entrance to the mRNA channel where it interferes with mRNA binding. We observe potent translation inhibition in the presence of Nsp1 in lysates from human cells. Based on the high-resolution structure of the 40S-Nsp1 complex, we identify residues of Nsp1 crucial for mediating translation inhibition. We further show that the full-length 5’ untranslated region of the genomic viral mRNA stimulates translation *in vitro*, suggesting that SARS-CoV-2 combines inhibition of translation by Nsp1 with efficient translation of the viral mRNA to achieve expression of viral genes^3^.

## Main

SARS-CoV-2 is the causing agent of the COVID-19 pandemic and belongs to the genus of beta-coronaviruses with an enveloped, positive sense and single-stranded genomic RNA^4^. Upon entering host cells, the viral genomic RNA is translated by the cellular protein synthesis machinery to produce a set of non-structural proteins (NSPs)^5^. NSPs render the cellular conditions favorable for viral infection and viral mRNA synthesis^6^. Coronaviruses have evolved specialized mechanisms to hijack the host gene expression machinery and employ cellular resources to regulate viral protein production. Such mechanisms are common for many viruses and include inhibition of host protein synthesis and endonucleolytic cleavage of host messenger RNAs (mRNAs)^1,7^. In cells infected with the closely related SARS-CoV, one of the most enigmatic viral proteins is the host shutoff factor Nsp1. Nsp1 is encoded by the gene closest to the 5’-end of the viral genome and is among the first proteins to be expressed after cell entry and infection to repress multiple steps of host protein expression^2,8,9,10^. Initial structural characterization of the isolated SARS-CoV Nsp1 protein revealed the structure of its N-terminal domain, whereas its C-terminal region was flexibly disordered^11^. Interestingly, SARS-CoV Nsp1 suppresses host innate immune functions, mainly by targeting type I interferon expression and antiviral signaling pathways^12^. Taken together, Nsp1 serves as a potential virulence factor for coronaviruses and represents an attractive target for live attenuated vaccine development^13,14^.

To provide molecular insights into the mechanism of Nsp1-mediated translation inhibition, we solved the structures of ribosomal complexes isolated from HEK293 lysates supplemented with recombinant purified Nsp1 as well as of an *in vitro* reconstituted 40S-Nsp1 complex using cryo-EM. We complement our findings by reporting *in vitro* translation inhibition in the presence of Nsp1 that is relieved after mutating key interacting residues. Furthermore, we show that the translation output of reporters containing full length viral 5’UTRs is greatly enhanced, which could explain how Nsp1 inhibits global translation while still translating sufficient amounts of viral mRNAs.

To elucidate the mechanism of how Nsp1 inhibits translation, we aimed to identify the structures of potential ribosomal complexes as binding targets. Previously, it has been suggested that Nsp1 mainly targets the ribosome at the translation initiation step^10^. Therefore, we treated lysed HEK293E with bacterially expressed and purified Nsp1 and loaded the cleared lysate on a sucrose gradient. Fractions containing ribosomal particles were then analyzed for the presence of Nsp1. Interestingly Nsp1 not only co-migrated with 40S particles, but also with 80S ribosomal complexes (Fig. 1a), suggesting that it interacts with a range of different ribosomal states. We then pooled all sucrose gradient fractions containing ribosomal complexes and investigated them using cryo-EM. This analysis revealed a 43S pre-initiation complex (PIC) encompassing the initiation factor eIF3 core, eIF1, the ternary complex comprising eIF2 and initiator tRNA_i_ with additional density in the mRNA entrance channel that could not be assigned to mRNA (Fig. 1b,c; Extended Data Fig. 1). To unambiguously attribute this extra density to Nsp1, we assessed whether Nsp1 binds purified ribosomal 40S subunits alone. *In vitro* binding assays using sucrose density centrifugation showed that Nsp1 associates with 40S ribosomal subunits since it co-pelleted with the 40S (Fig. 1d). However, Nsp1 did not interact with 60S subunits, suggesting that the interaction with 40S subunits is specific. Based on these results we assembled *in vitro* a 40S-Nsp1 complex and determined its structure at 2.8 Å resolution using cryo-EM (Extended Data Fig. 2). The molecular details revealed by these maps allowed us to identify the density as the C-terminal region of Nsp1 and build an atomic model. Docking of the model into the maps of the 43S PIC obtained from the HEK293E cell lysates clearly showed that the C-terminus of Nsp1 is also associated with the 43S PIC (Fig. 1b; Extended Data Fig. 3). As observed in the high-resolution structure of the 40S-Nsp1 complex, the C-terminal part of Nsp1 in the mRNA entrance channel (Fig. 1e) folds into two helices that interact with h18 of the 18S rRNA as well as proteins uS3 in the head and uS5 and eS30 in the body, respectively (Fig. 1f; Fig. 2a). In both complexes, Nsp1 binds in the mRNA entrance channel on the 40S subunit, where it would partially overlap with the fully accommodated mRNA. Consequently, mRNA was not observed due to Nsp1 binding.

**Figure 1:**
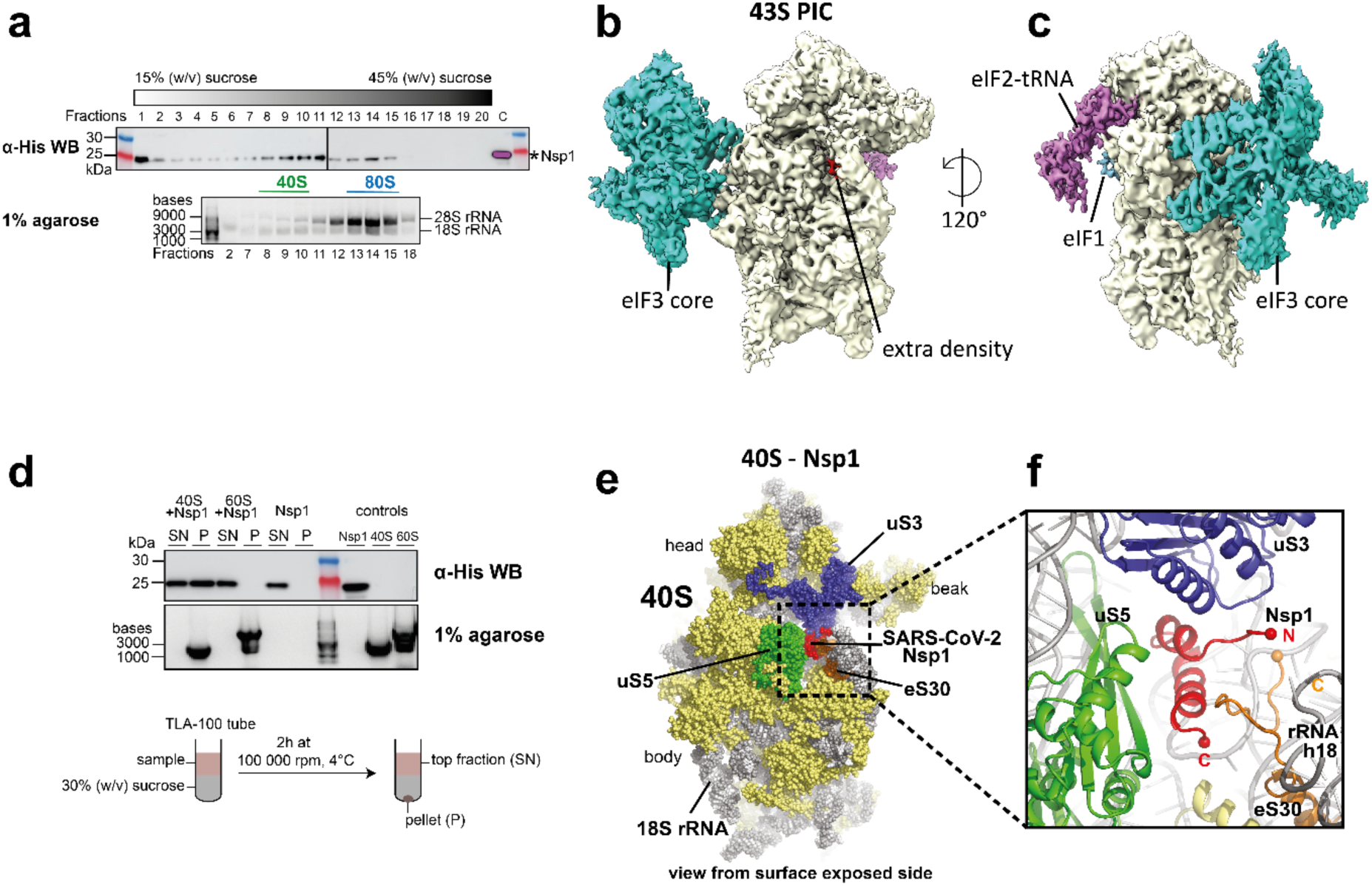
Structures of ribosomal complexes inhibited by SARS-CoV-2 Nsp1 solved by cryo-EM. **(a)** Sucrose gradient fractionation of HEK lysate supplemented with Nsp1. Nsp1 co-migrates with 40S and 80S ribosomal particles in a 15-45% (w/v) sucrose gradient. His_6_-tagged Nsp1 is visualized by Western blot using an α-His antibody, while the rRNA content in corresponding fractions is monitored on an agarose gel. All samples for the Western blot derive from the same experiment and the blots were processed in parallel. **(b-c)** Overview of Nsp1 (red) binding to a 43S PIC containing the core of initiation factor eIF3 (cyan), eIF1 (blue) and the eIF2-tRNA ternary complex (magenta). **(d)** In the *in vitro* binding assay, WT Nsp1 was added to 40S and 60S ribosomal SU and loaded on a 30% (w/v) sucrose cushion. Unbound proteins remained in the supernatant (SN), while bound Nsp1 co-pelleted with 40S (P). **(e)** Overview of Nsp1 binding to the small ribosomal subunit. Nsp1 (red) binds close to the mRNA entry site and contacts uS3 (blue) from the ribosomal 40S head as well as uS5 (green), the C-terminus of uS30 (orange) and h18 of the 18S rRNA (grey) of the 40S body. **(f)** Zoomed view of the area of Nsp1 binding as highlighted in (e).

**Figure 2:**
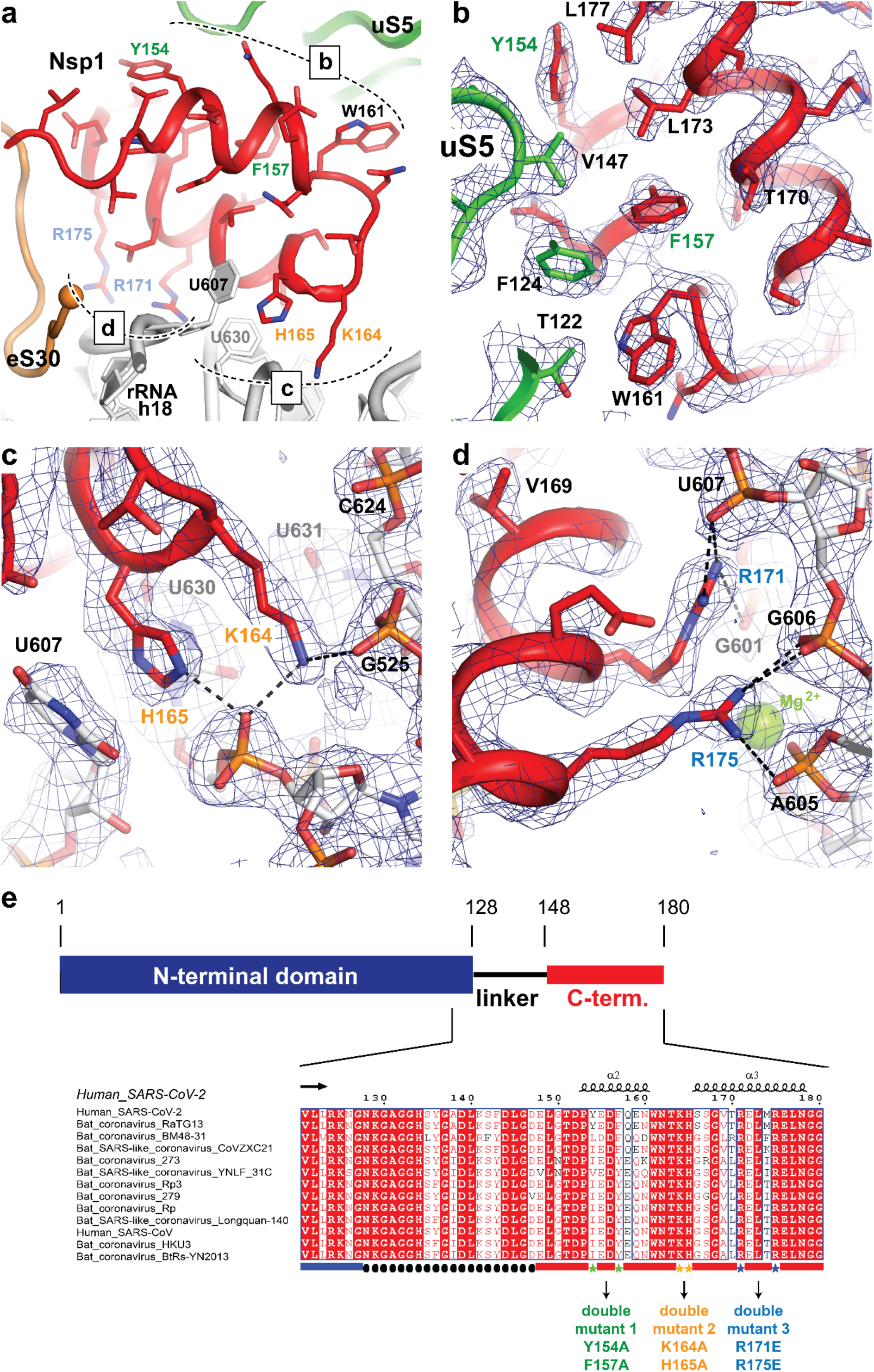
Binding of the C-terminal domain of SARS-CoV-2 Nsp1 to the 40S mRNA entry site is mediated by specific interactions *via* conserved residues. **(a)** Overview of Nsp1 binding to uS5 and the 18S rRNA. Nsp1 specifically interacts with uS5 through a hydrophobic patch and is tightly anchored to helix h18 of the 40S rRNA with a set of positively charged residues. **(b-d)** Detailed views of the specific interaction areas indicated in panel (b). The 2.8 Å experimental EM densities are shown as dark blue mesh and are contoured at 7.3s. Dashed lines indicate contacts between positively charged Nsp1 residues and the rRNA backbone within hydrogen bonding distance (< 3.8 Å). Residues mutated in this study are highlighted in red. **(e)** Alignment of the Nsp1 C-termini from human SARS-CoV, SARS-CoV-2 and closely related bat coronaviruses. Residues that mediate the interaction with the 40S mRNA entry site are highly conserved. Pairs of amino acids mutated for functional studies are highlighted. An overview of the Nsp1 domain arrangement including the flexible linker is depicted in the schematic.

The high-resolution reconstruction also revealed the network of molecular interactions between Nsp1 and the 40S subunit. The first C-terminal helix (residues 153-160) interacts with uS5 and uS3 through multiple hydrophobic side chains such as Y154, F157 and W161 (Fig. 2b). The two helices are connected by a short loop containing the KH motif that establishes stacking interactions with helix h18 of the 18S rRNA through U607 and U630 as well as backbone binding (Fig. 2c). The second helix (residues 166-178), localized in proximity of the eS30 C-terminus, interacts with the phosphate backbone of h18 *via* the two conserved arginines R171 and R175 (Fig. 2d).

An additional weak density at the head of the small subunit in the proximity of eS10 between h16 and uS3 was observed. This may correspond to the flexibly disposed N-terminal domain of Nsp1 considering the 20 amino acid long unstructured linker between the N-and C-terminal domains (Fig. 2e, Extended Data Fig. 4). However, we cannot exclude that this density corresponds to unassigned ribosomal protein segments in the vicinity as the C-terminal 65 amino acids of eS10 (head) or the N-terminal 60 amino acids of uS5 (body) could become better ordered in the context of Nsp1 binding. Thus, it occurs that Nsp1 is tightly bound to the 40S subunit through anchoring of its C-terminal helices to the mRNA channel, while the N-terminal domain can sample space in the radius of approximately 60 Å from its attachment point.

We further investigated how wild type (WT) Nsp1 affects translation of a *Renilla* Luciferase-encoding reporter mRNA (RLuc) in an S3 HeLa lysate *in vitro* translation system^15^ (Fig. 3d). WT Nsp1 was recombinantly expressed and purified and its effect on translation was tested by adding increasing amounts of the protein to HeLa cell lysates containing capped and polyadenylated RLuc mRNA control transcript. We observed a concentration-dependent inhibition of translation where almost full inhibition was reached at 1 µM of Nsp1 (Fig. 3b).

**Figure 3:**
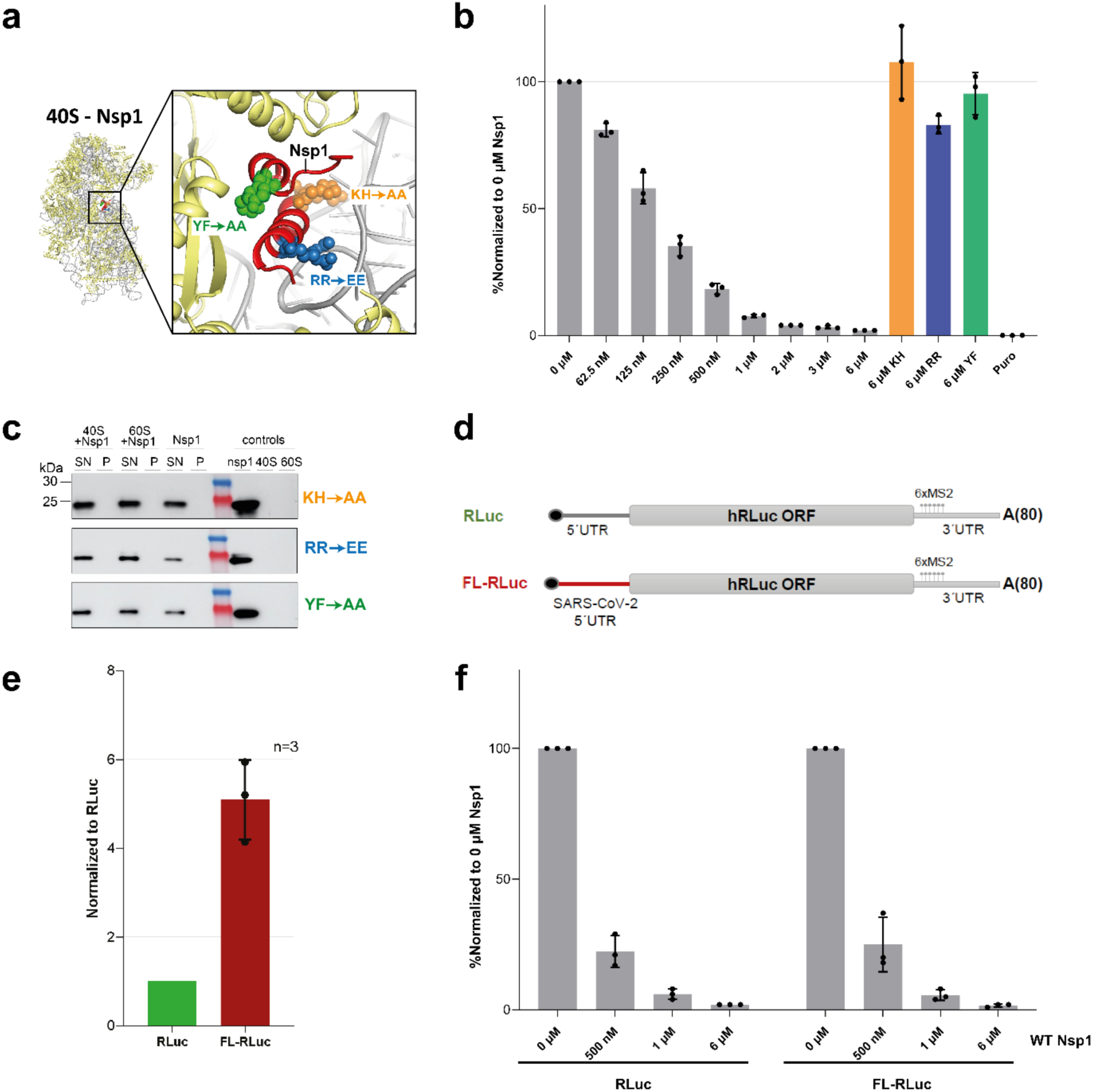
SARS-CoV-2 Nsp1 inhibits translation in HeLa cell lysates by binding to the 40S ribosomal subunit. **(a)** The C-terminal domain of Nsp1 binds to the mRNA entry site of 40S (red). Mutants in helix 2 (Y154A / F157A with spheres in green), in the short connecting loop (K164A / H165A with spheres in orange), and in helix 3 (R171E / R175E with spheres in blue) were generated. **(b)** Relative RLuc activity measurements of *in vitro* translation reactions normalized to the reaction in the absence of Nsp1 (0 μM). Mean values ± standard deviations of three biological replicates averaged after three measurements are shown, mean values of each biological replicate are indicated by dots. **(c)** *In vitro* binding assay, Nsp1 mutants were added to 40S and 60S ribosomal subunits and loaded on a 30% (w/v) sucrose cushion. SN: Supernatant; P: Pellet. **(d)** Schematic representation of the *in vitro* synthesized capped (black dot) and polyadenylated reporter mRNA coding for humanized Renilla Luciferase (hRLuc). **(e)** RLuc activity measurements of *in vitro* translation reactions using 40 fmol/µl of RLuc and FL-RLuc mRNA reporters, normalized to the readout of RLuc reporter mRNA. Mean values ± standard deviations of 3 biological replicates are shown, values of the individual measurements are indicated by dots. **(f)** Titration of WT-Nsp1 against RLuc and FL-RLuc reporter mRNAs. Relative RLuc activities were normalized to the untreated sample (0 μM). Mean values ± standard deviations of three biological replicates were averaged after three measurements are shown, mean values of each biological replicate are indicated by dots.

To dissect the contributions of the observed interactions between the 40S and Nsp1 on the inhibition of translation, we used our structural information to design several mutants targeting key amino acids in helix 1 (double mutant Y154A / F157A), the KH motif (double mutant K164A / H165A), and in helix 2 (double mutant R171E / R175E) as summarized in Fig. 3a. We also rationalized our mutations based on the high conservation of the Nsp1 C-terminus between SARS-CoV-2, SARS-CoV and closely related bat *Coronaviridae*, with sequence identities above 85% for the ORF1ab (encoding for polyproteins pp1a and pp1ab) and key amino acids highly conserved (Fig. 2e; Extended Data Fig. 4)^16^. Furthermore, the KH mutant had been described to abolish interaction with 40S in SARS-CoV^12^. In contrast to the WT protein, the three mutants did not affect translation of the RLuc control mRNA, even at concentrations of 6 µM (Fig. 3b). Consistently, the mutants lost their ability to bind 40S ribosomal subunits indicating that the C-terminal domain is primarily responsible for the affinity of Nsp1 for the ribosome (Fig. 3c). These results also agree with our structural findings where the C-terminal domain of Nsp1 is responsible for specific contacts with the ribosome, whereas the N-terminal domain is flexibly disposed. Interestingly, this mRNA binding inhibition mechanism may be unique to SARS-CoV-2 and closely related beta-coronaviruses, since the C-terminal region of Nsp1 is shorter in alpha-coronaviruses and is not highly conserved amongst other beta-coronaviruses including MERS-CoV, the latter being consistent with the observation that MERS-CoV Nsp1 does not bind the ribosome^17^.

Since translation of viral mRNA competes with translation of cellular mRNAs, the inhibitory effects of Nsp1 in the context of special features of the SARS-CoV-2 genomic RNA need to be considered. Therefore, we investigated the differences in the translation of reporter mRNAs with viral *vs.* cellular 5’UTRs and the relative inhibitory effect of Nsp1. Using the *in vitro* translation system described above, we compared the translation efficiency of RLuc reporters harboring the full-length 5’UTR of the SARS-CoV-2 genomic RNA (FL-RLuc) with the translation of equimolar amounts of a native RLuc reporter (Fig. 3d). We observed a significant five-fold increase in translation when the reporter mRNA included the viral 5’UTR, suggesting that the viral mRNA is more efficiently translated than host mRNAs (Fig. 3e). Nevertheless, titration of WT Nsp1 inhibited translation of both mRNAs, FL-RLuc and native RLuc, to the same extent (Fig. 3f).

These findings, as well as evidence from SARS-CoV^9^, indicate that Nsp1 acts as a general inhibitor of translation initiation. Our structural data suggest that SARS-CoV-2 Nsp1 inhibits translation by sterically occluding the entrance region of the mRNA channel and interfering with binding of cellular mRNAs (Fig. 4a,b). However, the question of how ribosomes in virus-infected cells are recruited to efficiently translate the viral mRNA remains open. Our results on the inhibitory mechanism of Nsp1 together with the translation-stimulating features of the viral mRNA provide a possible explanation. First, Nsp1 will act as a strong inhibitor of translation that tightly binds ribosomes and reduces the pool of available ribosomes that can engage in translation. Under ribosome limiting conditions, translation from more efficient viral mRNA is then likely to be favored (Fig. 4c). Therefore, the combination of an Nsp1-mediated general translation inhibition and an enhanced translation efficiency of viral transcripts appears to lead to an effective switch of translation from host cell mRNAs towards viral mRNAs.

**Figure 4:**
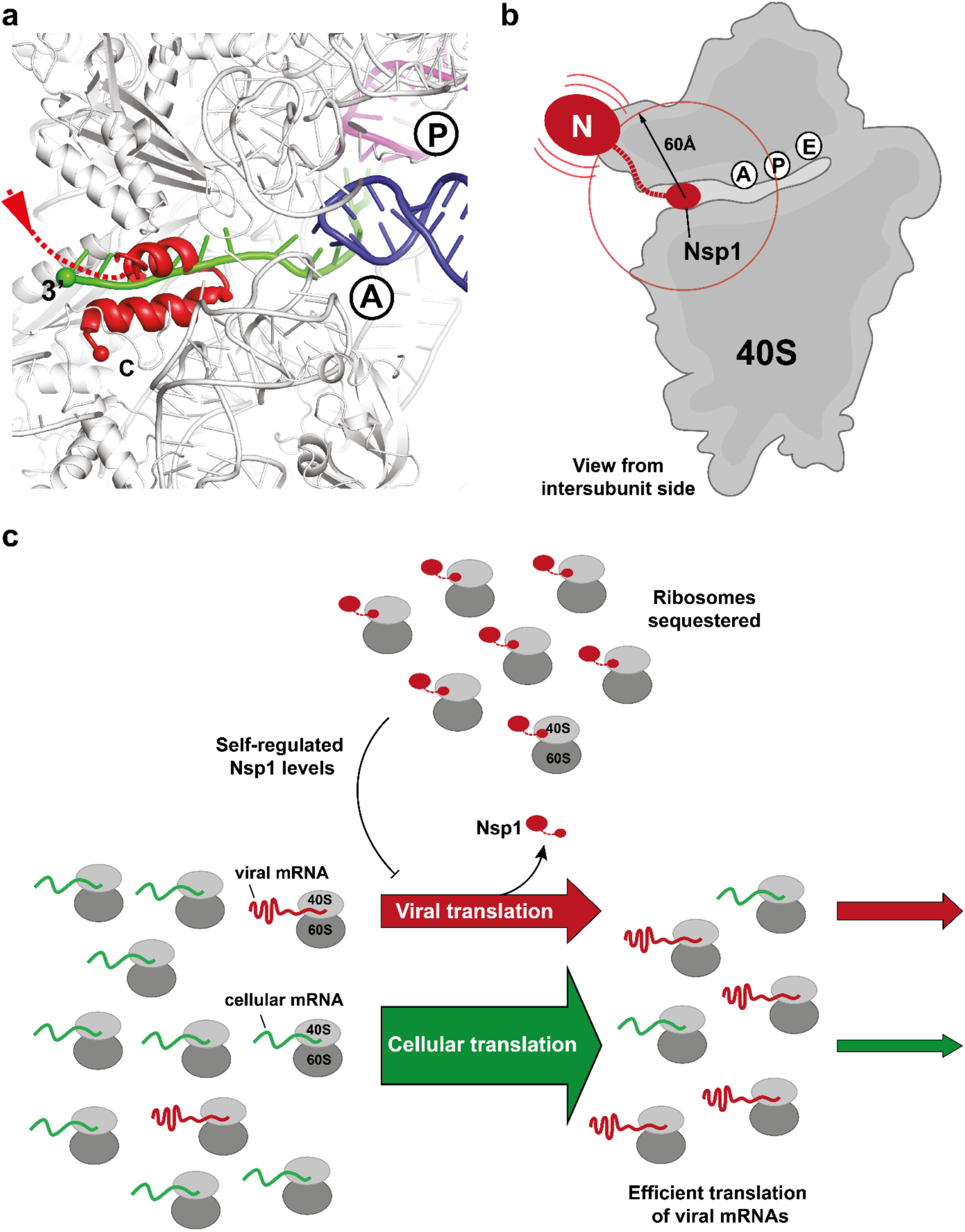
Binding of C-terminal domain of SARS-CoV-2 Nsp1 to ribosomal mRNA channel prevents classical mRNA binding by sterical hindrance. **(a)** Superposition of canonically bound mRNA (green), A-(blue) and P-site (purple) tRNAs (pdb 6HCJ) reveals that Nsp1 (red) prevents classical binding of the mRNA at the entry site due to blockage. **(b)** Nsp1 binds *via* its C-terminus in proximity of the 40S mRNA entry site. Due to the flexible linker, the N-terminal domain can sample an area of ∼60 Å around its attachment point (circle). **(c)** Model for translation inhibition by Nsp1. Upon viral infection and translation of viral genomic mRNA, Nsp1 acts as a translation inhibitor reducing the pool of ribosomes that can engage in translation. Under such ribosome-limiting conditions, viral mRNAs are translated with high efficiency.

Considering that Nsp1 can inhibit its own translation, the virus tunes cellular levels of Nsp1 exactly below the concentrations necessary to inhibit viral mRNA translation, but possibly enough to inhibit translation initiation from less efficiently recruited cellular mRNAs. Through this mechanism, we propose that Nsp1 would be able to inhibit global cellular translation particularly for mRNAs responsible for the host innate immune response, while the remaining ribosomes would still be able to translate viral mRNAs with high efficiency. During the course of viral infection, the effect of viral mRNAs on shifting protein synthesis machinery towards production of viral proteins would be increasingly strong since their levels are known to increase to 50% of total cellular RNAs^3^.

The identification of the C-terminal region of Nsp1 as the key domain for ribosome interactions that are essential for controlling cellular response to viral infections will be helpful in designing attenuated strains of SARS-CoV-2 for vaccine development. Furthermore, these results provide an excellent basis for structure-based experiments aimed at investigating Nsp1 function *in vivo* by using viral model systems.

## Methods

### Cloning, expression and purification of Nsp1 in *E.coli*

Plasmids encoding Nsp1 protein mutants were generated by site-directed mutagenesis using the primers 5’-GTT CAC GGG TCA CAC CGC TGC TAG CTG CGG TAT TCC AAT TTT CCT GAA AAT-3’, 5’-CGA GTT AGC CAC CAT TCA GTT CCT CCA TCA GTT CCT CGG TCA CAC CGC TGC TAT GTT TG −3’, 5’-TTT GGT ATT CCA ATT TTC CTG AGC ATC TTC AGC CGG ATC GGT GCC CAG TTC ATC G −3’ to yield KHAA, RREE and YFAA mutants, respectively. Nsp1 (WT and mutants) carrying an N-terminal His_6_-tag followed by a TEV cleavage site was expressed from a pET24a vector. The plasmid was transformed into *E. coli* BL21-CodonPlus (DE3)-RIPL and cells were grown in 2xYT medium at 30 °C. At an OD_600_ of 0.8, cultures were shifted to 18 °C and induced with IPTG added to a final concentration of 0.5 mM. After 16 h, cells were harvested by centrifugation, resuspended in lysis buffer (50 mM HEPES-KOH pH 7.6, 500 mM KCl, 5 mM MgCl_2_, 40 mM imidazole, 10% (w/v) glycerol, 0.5 mM TCEP and protease inhibitors) and lysed using a cell disrupter (Constant Systems Ltd). The lysate was cleared by centrifugation for 45 min at 48.000 xg and loaded onto a HisTrap FF 5-ml column (GE Healthcare). Eluted proteins were incubated with TEV protease at 4 °C overnight and both the His_6_-tag, uncleaved Nsp1 and the His_6_-tagged TEV protease were removed on the HisTrap FF 5-ml column. The sample was further purified via size-exclusion chromatography on a HiLoad 16/60 Superdex75 (GE Healthcare), buffer exchanging the sample to the storage buffer (40 mM HEPES-KOH pH 7.6, 200 mM KCl, 40 mM MgCl_2_, 10% (w/v) glycerol, 1 mM TCEP). Fractions containing Nsp1 were pooled, concentrated in an Amicon Ultra-15 centrifugal filter (10-kDa MW cut-off), flash-frozen in liquid nitrogen, and stored until further use at −80 °C.

### Preparation of human ribosomal subunits

Human ribosomal subunits were purified as described^18^, and final samples were flash-frozen in liquid nitrogen at a concentration of 1 mg/ml (OD_600_ of 10) and stored at −80 °C.

### Sucrose pelleting binding assay

To verify Nsp1-40S complex formation, we performed binding assays using sucrose density centrifugation. Thawed human 40S and 60S ribosomal subunits were adjusted to a final concentration of 0.3 μM in a 100 μl reaction (complex binding buffer: 20 mM HEPES pH 7.6, 5 mM MgCl_2_, 100 mM KCl, 2 mM DTT) and mixed with a 5x molar excess of His_6_-Nsp1 WT/mutant. The assembled complexes were incubated for 5 min at 30 °C and for 10 min on ice before they were loaded on a 30% (w/v) sucrose cushion in TLA-100 tubes (Beckman Coulter). The sucrose cushions were centrifuged for 2 h at 390,880 xg and 4 °C. After removing the supernatant, the pellet was resuspended in 20 μl of complex binding buffer. Samples were analyzed on bleach agarose gels (0.06% bleach, 1% (w/v) agarose) for visualization of the RNA and on WB (anti-His antibody, Clontech) for visualization of His_6_-tagged Nsp1.

### Preparation of viral 5’-UTR mRNA and reporter RLuc mRNA

#### Plasmids

MS2-containing mRNA reporters (p200-6xMS2) were derived from the pCRII-hRLuc-200bp 3’UTR construct as described before^15^. Amplification of the vector using the primers 5’-AAT AAG AGC TCC TGC CTC GAG CTT CCT CATC-3’ and 5’-AAT AAC ATA TGG TGA TGC TAT TGC TTT ATT TGT AAC-3’ was followed by restriction digestion with SacI and NdeI and ligation with a 6xMS2-containing insert that was PCR-amplified using the primers 5’-AAT AAC ATA TGG TTC CCT AAG TCC AAC TAC CAA A-3’ and 5’-AAT AAA GAG CTC CCA GAG GTT GAT TGT CGA CC-3’ and that had been treated with the same enzymes. The 223 nt-long 5’UTR of SARS-CoV genomic mRNA sequence was subcloned to replace the RLuc 5’ UTR by fusion PCR using primers TCTGCAGAATTCGCCCTTCATG and GCCCTATAGTGAGTCGTATTACAATTCACT for vector amplification and the pair GACTCACTATAGGGCAACTTTAAAATCTGTGTGGCTGTCACT and GGCGAATTCTGCAGACTTACCTTTCGGTCACACCCG for amplification of the 5’UTR fragment using 5’UTR-eGFP cloned in pUC19 vector as a template, which was designed to possess the SARS-CoV-2 5’UTR sequence in front of the eGFP coding sequence.

#### *In vitro* transcription of reporter mRNAs

Preparation of *in vitro* transcribed mRNAs was performed as described^15^. Namely, linearized pCRII vectors encoding the desired reporter mRNA downstream of a T7 promoter were mixed to yield an *in vitro* transcription reaction in 1x Transcription Buffer (Thermo Fisher Scientific) at a final concentration of 20-30 ng/µl. This mixture further contained 1 mM of each ribonucleotide (rNTPs, Thermo Fisher Scientific), 1 u/µl Murine RNase inhibitor (Vazyme), 0.001 u/µl Pyrophosphatase (Thermo Fisher Scientific) and 5% (v/v) T7-RNA-polymerase (custom-made). The reaction was incubated at 37 °C for 1 h and then an equal quantity of T7-RNA polymerase was added for another 30 min. The mixture was then supplemented with TURBO DNase (Thermo Fisher Scientific) to a final concentration of 0.14 u/µl and incubated at 37 °C for 30 min. The transcribed mRNA was purified from the reaction using an acidic phenol-chloroform-isoamylalcohol (P.C.I.). The product was dissolved in disodium citrate buffer, pH 6.5 and quality was assessed by agarose gel electrophoresis.

Prior to capping, the RNA was incubated at 65 °C for 5 min and supplemented accordingly to yield a reaction consisting of 300 ng/µl RNA, 0.5 mM guanosine triphosphate (GTP, New England Biolabs), 0.1 mM S-adenosylmethionine (SAM, New England Biolabs), 1 u/µl Murine RNase inhibitor (Vazyme), 0.5 u/µl vaccinia capping enzyme (VCE, New England Biolabs) in 1x Capping buffer (New England Biolabs). The capping reaction was carried out at 37 °C for 1 h and quenched by the addition of acidic P.C.I., followed by RNA purification. Finally, the integrity of the capped mRNAs was verified by agarose gel electrophoresis.

### Preparation of HeLa translation-competent lysates

HeLa S3 lysates were prepared similarly as described before^15^. Briefly, lysates were prepared from S3 HeLa cell cultures grown to a cell density ranging from 1-2×10^6^ cells/ml. Cells were pelleted (200 *g*, 4 °C for 5 min) and washed two times with cold PBS pH 7.4 and finally resuspended in ice-cold hypotonic lysis buffer [10 mM HEPES pH 7.3, 10 mM K-acetate, 500 μM Mg-acetate, 5 mM DTT and 1x protease inhibitor cocktail (biotool.com)] at a final concentration of 2×10^8^ cells/ml. The suspension was incubated on ice for 10 min and cells were lysed by dual centrifugation (500 rpm, −5 °C, 4 min) using Zentrimix 380R (Hettich) with a 3206 rotor and 3209 adapters. The lysis process was monitored by trypan stain. The lysate was centrifuged at 13’000 x*g*, 4 °C for 10 min and the supernatant was aliquoted, snap frozen and stored at −80 °C.

### *In vitro* translation assays

*In vitro* translation reactions were performed similarly as described before^15^. Briefly, 400 μl of recombinant proteins were dialyzed overnight in 30 mM NaCl, 5 mM Hepes pH 7.3 at 4 °C using Slide-A-Lyzer MINI Dialysis devices with a 3.5K MWCO (Thermo Scientific, 88400), and the protein concentration was calculated using Nanodrop. In parallel, equal volumes of recombinant protein storage buffer were dialyzed and used as negative control (0 μM condition) and to maintain the same concentration of dialyzed storage buffer in translation mixtures. S3 lysate corresponding to 1.11×10^6^ cell equivalents was used at a concentration of 8.88 × 10^7^ cell equivalents/ml. The reaction was supplemented to a final concentration of 15 mM HEPES, pH 7.3, 0.3 mM MgCl_2_, 24 mM KCl, 28 mM K-acetate, 6 mM creatine phosphate (Roche), 102 ng/µl creatine kinase (Roche), 0.4 mM amino acid mixture (Promega) and 1 u/µl NxGen RNase inhibitor (Lucigen). Control reactions contained 320 µg/ml puromycin (Santa Cruz Biotechnology) and all reactions were complemented with an equal volume of dialyzed protein purification buffer. In all reactions where recombinant Nsp1 was used, the lysate was pre-incubated with Nsp1 at 4 °C for 30 minutes. Before addition of reporter mRNAs, the mixtures were incubated at 33 °C for 5 min. *In vitro* transcribed and capped mRNAs were incubated for 5 min at 65 °C,15 min at RT and cooled down on ice. Reporter mRNAs were added to the translation reactions at a final concentration of 40 fmol/μl. The translation reaction was performed at 33 °C for 50 min. To monitor the protein synthesis output, samples were put on ice and mixed with 50 μL 1x Renilla-Glo substrate (Promega) in Renilla-Glo (Promega) assay buffer on a white bottom 96 well plate. The plate was incubated at 30 °C for 10 min and the luminescence signal was measured three times using the TECAN infinite M100 Pro plate reader and plotted on GraphPad after performing three independent biological replicates.

### HEK extracts and sucrose gradient analysis

HEK293E cell lysates were supplemented with Nsp1 to purify native-like Nsp1-inhibited translation complexes. For this, frozen HEK293E cells were thawed and resuspended in 2x excess of lysis buffer (25 mM HEPES-KOH pH 7.6, 5 mM MgCl_2_, 50 mM KCl, cOmplete protease inhibitor cocktail (Roche), 1 mM PMSF, 0.2 U/µl RiboLock). For cell lysis, cells were transferred to a Dounce homogenizer (tight) and lysed with 12 strokes. After adding Triton X-100 to a final concentration of 0.1%, the lysate was incubated under rotation for 30 min at 4 °C and cleared for 10 min in an MLA-80 rotor (Beckman Coulter) at 11,500 xg and 4 °C. The cleared HEK293E lysate was treated with 100 µg/ml cycloheximide and 1 mM GMP-PNP for 5 min at 30 °C. Nsp1 (for cryo-EM) or His_6_-Nsp1 (for Western blot) was added to a final concentration of 2 µM, and the extracts were incubated for additional 5 min at 30 °C before they were loaded onto 15% - 45% (w/v) sucrose gradients (20 mM HEPES-KOH pH 7.6, 100 mM KOAc, 5 mM MgCl_2_, 1 mM DTT). Gradients were centrifuged in a SW 32.1 Ti rotor (Beckman) at 79,500 xg for 15 h at 4 °C and manually fractioned with a syringe. Fractions containing ribosomal particles were pooled and concentrated in an Amicon Ultra-15 centrifugal filter (100-kDa MW cut-off). For biochemical analyses, the same gradients were prepared, with the exception of using His_6_-tagged Nsp1 instead of the TEV-cleaved protein. Fractions were precipitated with trichloroacetic acid (TCA) and subjected to Western blot analysis (anti-His antibody, Clontech). Additionally, before precipitation, samples were taken for analysis on agarose gels (0.06% bleach, 1% (w/v) agarose).

### Cryo-EM sample preparation and data collection

Quantifoil R2/2 holey carbon copper grids (Quantifoil Micro Tool) were prepared by first applying an additional thin layer of continuous carbon and then glow-discharging them for 15 sec at 15 mA using an easiGlow Discharge cleaning system (PELCO). For the *in vitro* binding experiment, purified Nsp1 was first mixed with 40S in molar ratio of 10:1. For the HEK lysate sample, sucrose peak fractions containing ribosomes were collected, buffer exchanged and concentrated. Then, 4 µL samples at concentrations of 80 – 100 nM of the 40S-nsp1 or ribosomes from the HEK cell lysate were applied to the grids, which were then blotted for approximately 8 s and immediately plunged in 1:2 ethane:propane (Carbagas) at liquid nitrogen temperature using a Vitrobot (Thermo Fisher Scientific). The Vitrobot chamber was kept at 4 °C and 100% humidity during the whole procedure.

For each sample, one grid was selected for data collection using a Titan Krios cryo-transmission electron microscope (Thermo Fisher Scientific) operating at 300 kV and equipped with either a Falcon3EC camera (Thermo Fisher Scientific) in integration mode or a K3 camera (Gatan), which was run in counting and super-resolution mode, mounted to a GIF Quantum LS operated with an energy filter slit width of 20 eV. The Falcon3EC datasets were collected at a nominal magnification of 75’000 x (pixel size of 1.08 Å/pixel), while for the K3 datasets a nominal magnification of 81’000x was used (physical pixel size of 1.08 Å/pixel, which corresponds to a super-resolution pixel size of 0.54 Å/pixel). For counting mode, illumination conditions were adjusted to an exposure rate of 24 e^-^/pixel/second. Micrographs were recorded as movie stacks at an electron dose of ∼60 e^-^/Å^2^ applied over 40 frames. For both datasets, the defocus was varied from approximately −1 to −3 μm.

### Cryo-EM data processing

The stacks of frames were first aligned to correct for motion during exposure, dose-weighted and gain-corrected using MotionCor2^19^. The super-resolution micrographs collected with the K3 camera were additionally binned 2 times during the MotionCor2 procedure. The contrast transfer function of the motion-corrected and dose-weighted micrographs were then estimated using GCTF^20^.

Micrographs (10’104 for the *in vitro* binding experiment, 16’887 for the HEK cell extract) were carefully inspected based on CTF estimations for drift and ice quality. Particle images of ribosomes were picked (2’078’577 for the *in vitro* binding experiment, 1’080’818 for the HEK cell extract) in Relion3.1 using a Laplacian-of-Gaussian filter-based method^21^. The picked particle images were then subjected to a reference-free 2D classification in RELION/cryoSPARC2, and the particles were selected from the 2D class-averages (1’718’196 particles for the *in vitro* binding experiment, 619’890 for the HEK cell extract). For the *in vitro* binding experiment, the particles were then classified in 3D using a human 40S re-initiation complex (EMD-3770^18^) that was low-pass filtered to 60 Å to select for good 40S classes, followed by refinement using Relion3.1^22^. Further processing was done with Relion3.1 and cryoSPARC2^23^ according to the scheme shown in Extended Data Fig. 2. Transformation of particle information between the two programs was done using PyEM script (Asarnow, D., Palovcak, E., Cheng, Y. UCSF pyem v0.5. Zenodo https://doi.org/10.5281/zenodo.3576630 (2019)). In short, the particle set was first cleaned from the preferentially oriented particles based on their orientation parameters, which reduced the particle set to 700’459 particles. Those particles were then further classified for their quality and for the presence of Nsp1 using a focused 3D classification approach. The final set of particle images was refined using a global 3D refinement. To further improve the local resolution of the 40S-Nsp1 complex, masks around the 40S head and body were generated using UCSF Chimera^24^ by creating a mask which was extended by 10 Å around a fitted model of the 40S subunit. Those masks were used for a multi-body refinement in Relion3.1^25^. Finally, the two focused maps were combined to generate a composite 3D map of the entire *in vitro* reconstituted 40S-Nsp1 complex.

For the HEK cell extract, after 2D classifications, *ab initio* reconstruction was performed in cryoSPARC2^23^, and the determined volumes were used as starting references for a heterogeneous refinement in cryoSPARC2 (Extended Data Fig. 1). The 80’101 particle images corresponding to the 40S ribosomal subunit were selected for a further round of heterogeneous refinement in cryoSPARC2, which resolved a density corresponding to initiation factor eIF3 in a fraction of the particles. To improve the occupancy of eIF3, particle images belonging to the 40S subunit class were then subjected to a focused 3D classification in Relion3.1 using a circular mask on the eIF3 region. The 3D class depicting the best density for eIF3 was selected (8’000 particle images) and was then used for a global 3D refinement. To further improve the resolution of the Nsp1-bound region, a focused refinement was done using a mask on the body of the 40S subunit.

### Structure building and refinement

For building of the 40S-Nsp1 complex, the head and body of PDB 5oa3^18^ were docked as rigid bodies into the 2.8 Å head and body maps that were obtained by focused classification (Extended Data Fig. 2). The structures were adjusted manually into the high-resolution maps using COOT^26^, and the C-terminus of Nsp1 (residues 148-180), which was well-resolved in the map of the 40S body, was built *de novo*. The coordinates were subjected to 5 cycles of real space refinement using PHENIX 1.18^27^. To stabilize the refinement in less well-resolved peripheral areas, protein secondary structure and Ramachandran as well as RNA base pair restrains were applied. Remaining discrepancies between models and maps as well as missing Mg^2+^ ions were detected using real space difference maps, and after model completion the coordinates were refined for two additional cycles. The resulting final models have excellent geometries and correlations between the maps and models (Table 1, Extended Data Fig. 2). The structures were validated using MOLPROBITY^28^ and by comparison of the model vs. map FSCs at values of 0.5, which coincided well with the FSCs between the half-sets of the EM reconstruction using the FSC=1.43 criterion (Extended Data Fig. 2).

To assemble the full 40S-Nsp1 complex, both refined structures were docked into a 2.8 Å chimeric map comprising the complete 40S-Nsp1. After readjustment of the head-to-body connections, the complete model was subjected to two additional rounds of real space refinement as described above.

The 5.9 Å and 4.3 Å maps of the Nsp1-43S PIC shown in Extended Data Fig. 1 and 3 were aligned onto the 2.8 Å 40S-Nsp1 body map of the *in vitro* reconstituted complex in UCSF Chimera, into which the atomic model had been built. For general interpretation of 43S PIC, the refined models of the 40S head and body determined for the in-vitro 40S-Nsp1 complex were docked as rigid bodies in UCSF Chimera^28^. Initiation factors IF2 and IF3 were taken from pdb 6YAM^29^ and docked similarly. A homology model for the missing eIF2β subunit was obtained using PHYRE2^30^ and pdb 3JAP^31^ as a template, and the density of IF1 was interpreted using pdb 2IF1^32^.

## Data availability

The high-resolution cryo-EM maps of the complete 40S-Nsp1 complex, the 40S body and the 40S head-Nsp1 have been deposited in the Electron Microscopy Data Bank as EMD-11320, EMD-11321 and EMD-11322, respectively, while the corresponding models are in the Protein Data Bank as PDB IDs 6ZOJ, 6ZOK, 6ZOL. Additionally, the 5.9 Å resolution map for the 43S PIC and the 4.3 Å map focused on the body of the small ribosomal subunit in the 43S were deposited in the Electron Microscopy Data Bank as EMD-11323 and EMD-11324.

## Acknowledgements

We thank the ETH Scientific center for optical and electron microscopy (ScopeM) and the CryoEM Knowledge hub (CEMK), in particular D. Böhringer, for technical support and the opportunity to continue our work in spite of the ETH lockdown due to the COVID-19 pandemic. We thank the Functional Genomics Center Zurich (FGCZ) for the help with mass-spectrometry. The authors would like to thank their teams for the support in the lab, and especially to M. Jia, P. Bhatt and D. Yudin for creating a productive working atmosphere.

## Funding

This work was supported by grants of NB, OM and VT from the Swiss National Science Foundation (SNSF; grant numbers 173085, 182341 and 182831), the National Center of Competence in Research (NCCR) on RNA and Disease funded by the SNSF, the ETH Research Grant ETH-23 18-2 and a Ph.D. fellowship by Böhringer Ingelheim Fonds to KS.

## Authors’ contributions

NB and KS initiated the project and designed the experiments. KS expressed proteins, together with BE, and prepared samples for cryo-EM. KS, AJ and AS prepared grids, carried out data collection and processing. EDK and OM designed translation experiments, EDK and LG were involved in cloning and EDK performed *in vitro* translation reactions, with the help of LG. KS and BE performed sucrose binding assays.

ML was involved in structure modelling and refinement as well as in figure preparation. NB and KS coordinated the project. All authors contributed to the final version of the manuscript.

## Conflicts of interest

The authors declare no competing interests.

## Supplementary figures

**Extended Data Figure 1:**
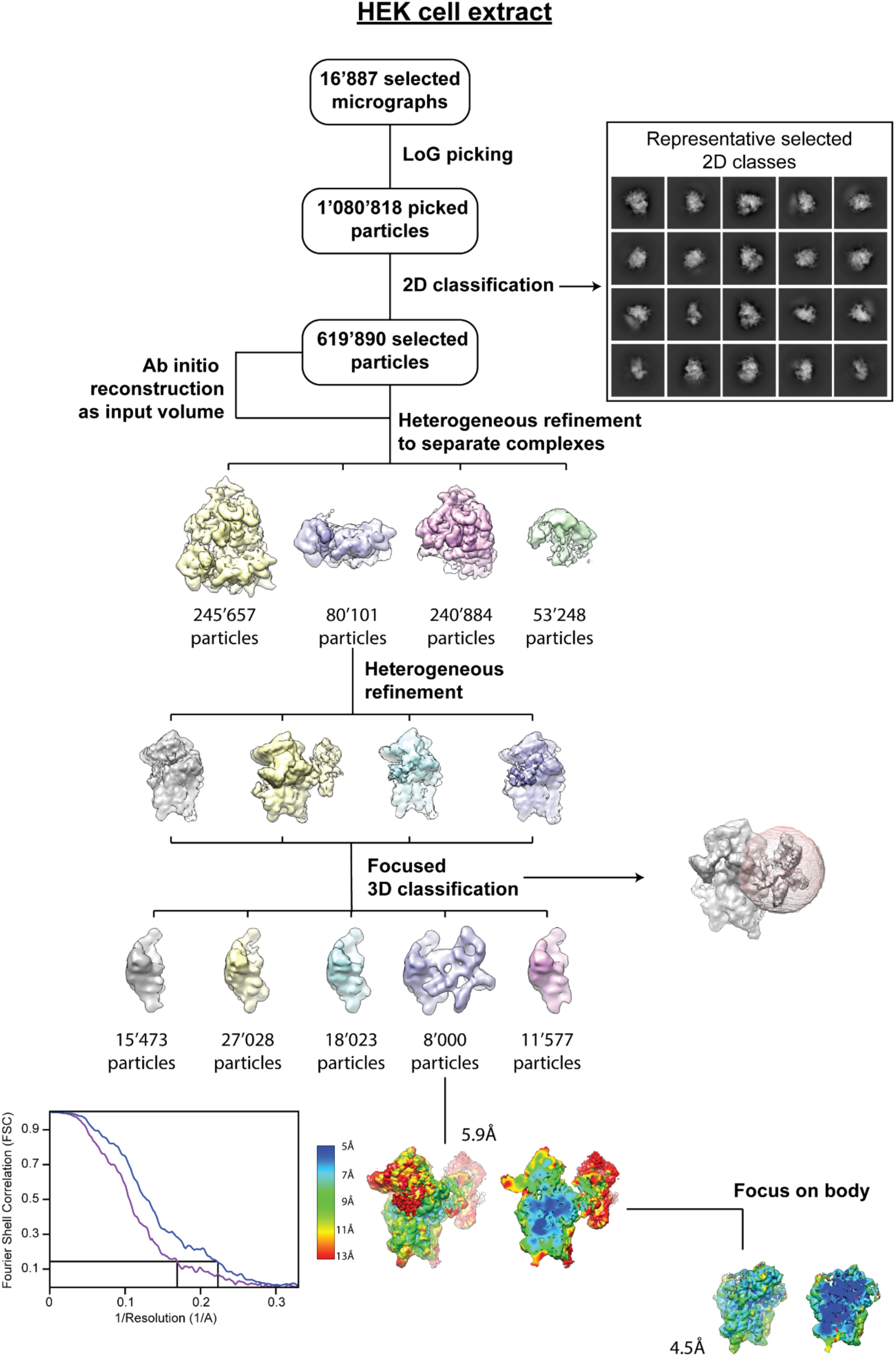
Data processing of the HEK cell extract cryo-EM dataset. Scheme for the processing of the HEK cell extract sample. Local resolution estimates are plotted as heat map on the final volume accompanied with a slice through the volume. The half map vs. half map FSC curves are shown for the overall refinement (purple) and the refinement focused on the body (blue).

**Extended Data Figure 2:**
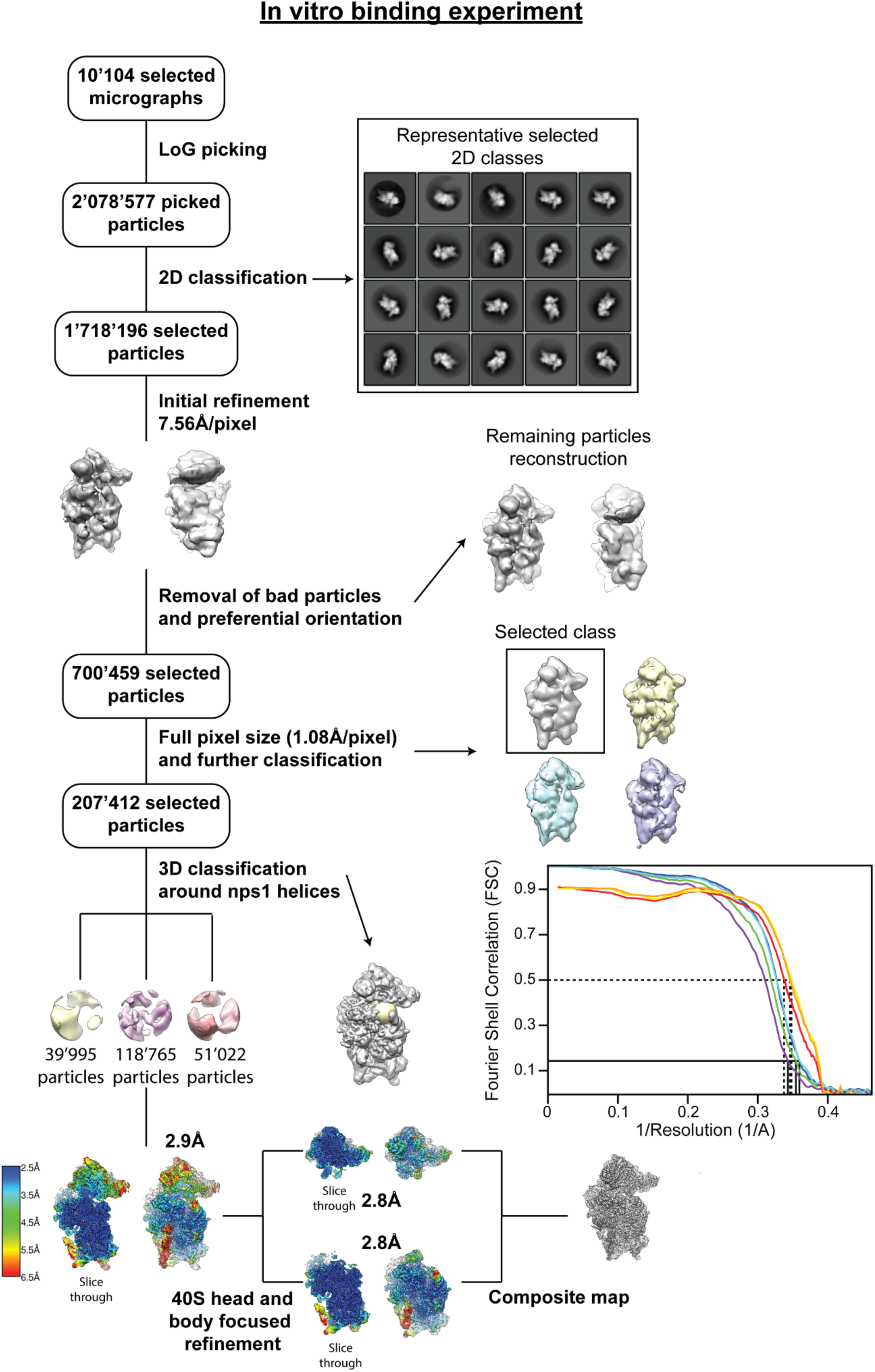
Data processing of the *in vitro* reconstituted 40S-Nsp1 cryo-EM dataset. Scheme of the processing steps performed for the sample of the *in vitro* binding experiment. The local resolution distribution is plotted on the final volumes as heat map, together with additional slices through the volumes. The half map vs. half map FSC curves are plotted for the overall refinement (purple), the refinement focused on the body (blue), on the head (cyan), as well as for the composite map (green). The map vs. model FSCs are plotted for the body (yellow) and the head (orange) in their respective focused maps, as well as for the full 40S in the composite map (red).

**Extended Data Figure 3:**
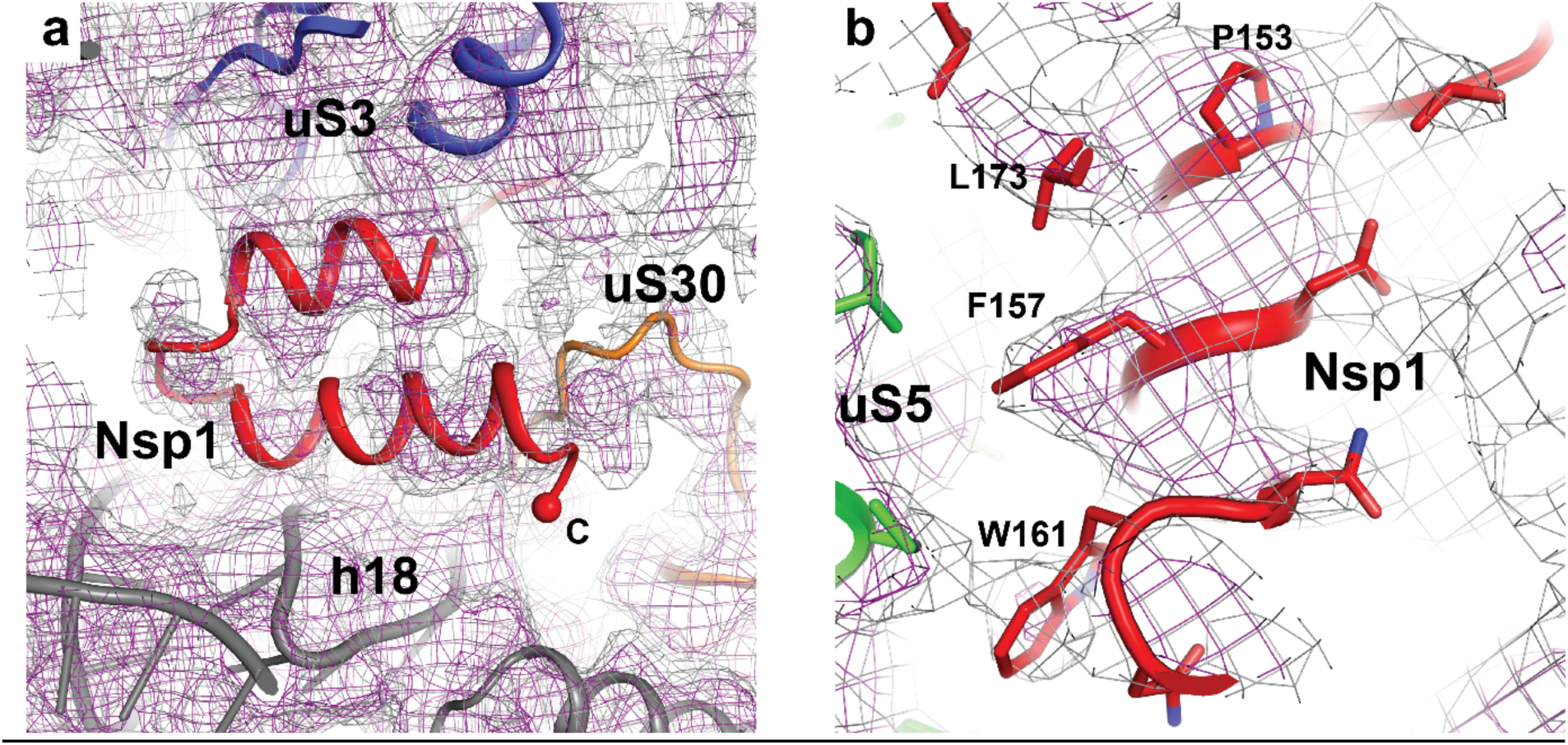
Binding of Nsp-1 to the 43S initiation complex. **(a)** Additional EM density is present in the 40S mRNA entry site. Docking of the high-resolution 40S-Nsp1 structure into the 4.3 Å map focused on the body reveals an excellent fit of the two C-terminal Nsp1 helices. **(b)** For several bulky side chains mediating the hydrophobic contact with uS5 of the 40S body, side chain densities can be recognized.

**Extended Data Figure 4:**
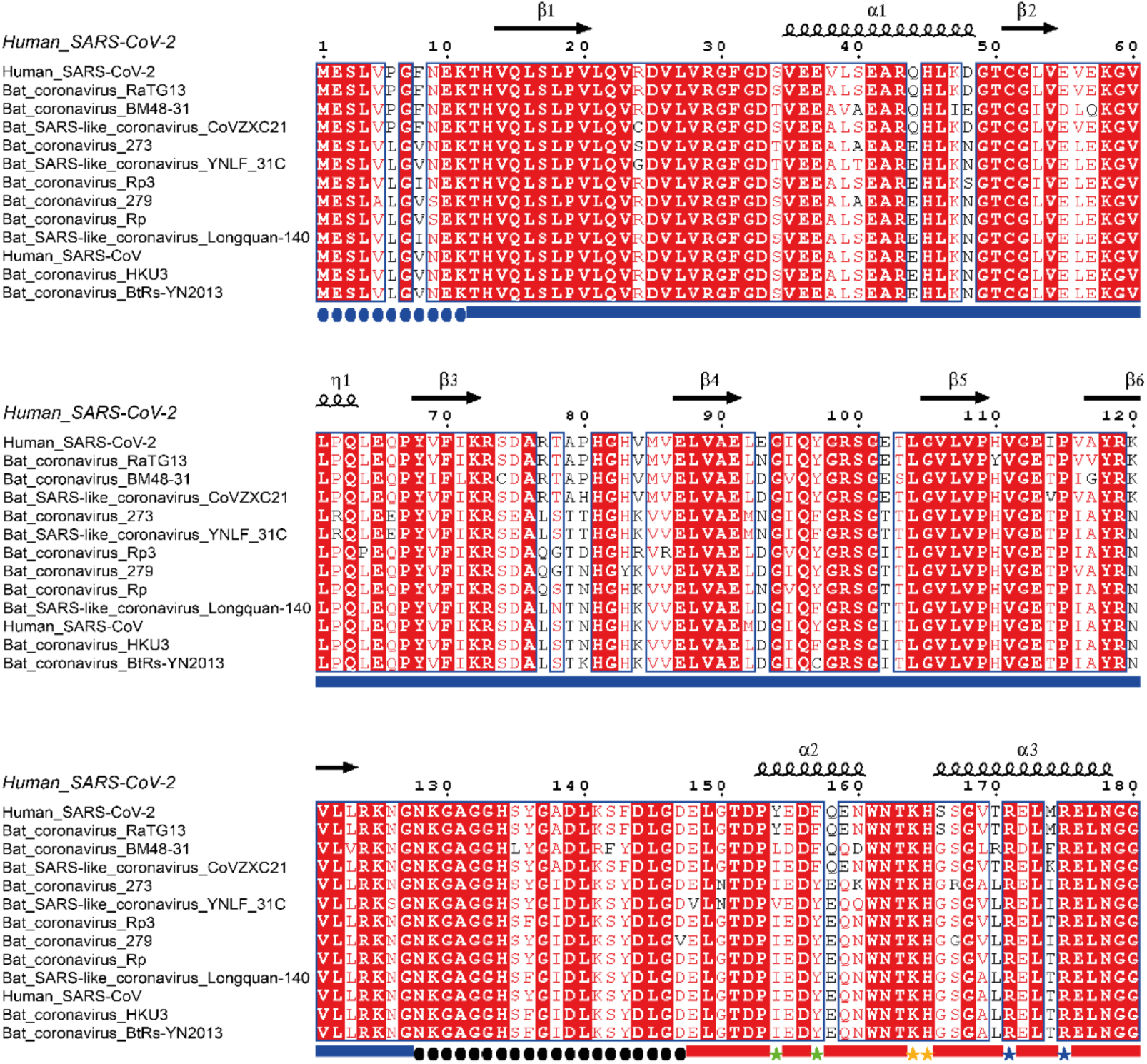
Alignment of full-length Nsp1 sequences from selected beta-coronaviruses. Nsp1 sequences of human SARS-CoV, human SARS-CoV-2 and SARS-related bat coronaviruses^16^ were obtained from the UNIPROT (www.uniprot.org) and GenBank (www.ncbi.nlm.nih.gov/genbank) databases. The sequences were aligned using Clustal Omega (www.ebi.ac.uk/Tools/msa/clustalo). The alignment was visualized with ESPript^33^. Note that sequences of MERS Nsp1 and other human coronaviruses were not included in the alignment due to lack of sequence homology. For displaying the secondary structure, the atomic coordinates of the SARS-CoV Nsp1 N-terminus (pdb 2HSX)^11^ and of the Nsp1 C-terminus (this publication) were combined. Regions of known structure are highlighted with blue (N-terminal domain) and red (C-terminal domain) bars. Unresolved regions are indicated by dotted lines, including the ∼60 Å unstructured linker (black) and the N-terminus (blue). Double mutations analyzed in this study are shown as asterisks in different colors.

**Extended Data Figure 5:**
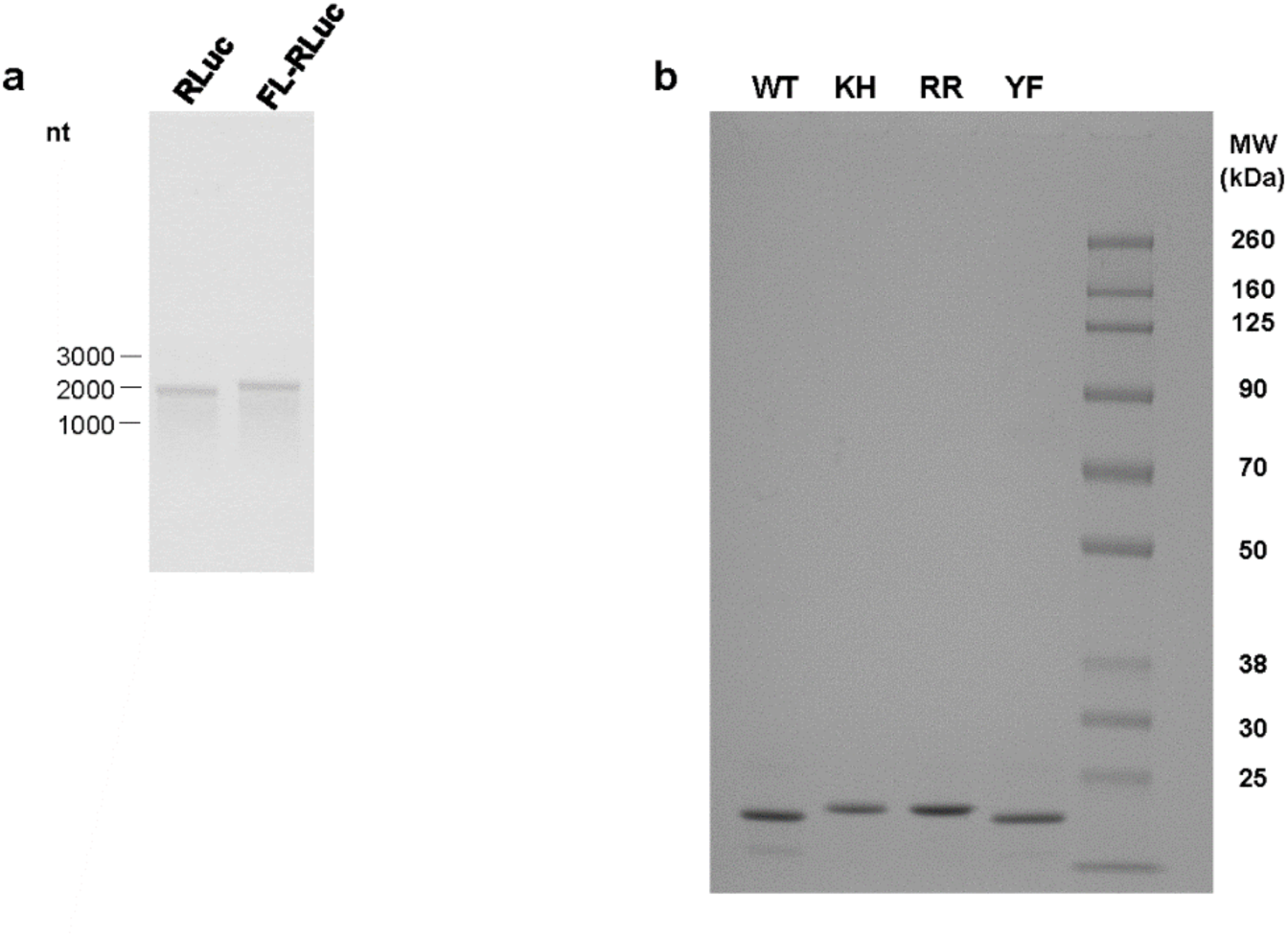
Additional components of *in vitro* translation reaction. **(a)** 1% agarose gel electrophoresis of the *in vitro* transcribed RLuc and FL-RLuc reporters used for the *in vitro* translation assays. **(b)** Input samples of the *in vitro* translation assay described in Fig. 3b. 2 μL of dialyzed Nsp1 protein samples at a concentration of 17.9 μM were analyzed on a 4-12% SDS-PAGE and stained by Imperial protein staining (Thermo Fisher Scientific).

**Extended Table 1:** Cryo-EM data collection, map refinement, model refinement and validation statistics of the human 40S in complex with Nsp1.

| Structure | 40S head | 40S body – Nsp1 | complete 40S - Nsp1 |
| --- | --- | --- | --- |
| EMDB accession number / PDB code | #### / #### | #### / #### | #### / #### |
| Data collection |  |  |  |
| Microscope | TFS Titan Krios G3i |  |  |
| Detector | Gatan K3 direct electron detector |  |  |
| Voltage (keV) | 300 |  |  |
| Electron exposure (e<sup>-</sup>/Å²) | 60 |  |  |
| Pixel size (Å) | 1.08 (super-resolution pixel at 0.54Å/pixel) |  |  |
| Magnification | 81'000x (nominal) |  |  |
| Defocus range (µm) | 1-3 |  |  |
| Automation software | EPU2 |  |  |
| Selected micrographs | 10'104 |  |  |
| EM Reconstruction |  |  |  |
| Initial particle images (no.) | 2'078'577 | 2'078'577 | 2'078'577 |
| Final particles (no.) | 118'765 | 118'765 | 118'765 |
| Resolution at FSC=0.143 (Å) | 2.8 | 2.8 | 2.9 |
| Sharpening B-factor (Å²) | −62.2 | −62.8 | −70 |
| Coordinate real space refinement (PHENIX version 1.18.2-3874) |  |  |  |
| Unit cell | P1 | P1 | P1 |
| a, b, c (Å) | 181.44, 140.40, 114.48 | 181.44, 189.00, 201.96 | 206.28, 231.12, 248.40 |
| α = β = γ (°) | 90 | 90 | 90 |
| CC<sub>mask</sub> | 0.87 | 0.86 | 0.85 |
| High resolution limit for refinement | 2.8 | 2.8 | 2.8 |
| Resolution according to model vs. map FSC=0.5 (masked) criterion (Å) | 2.8 | 2.8 | 2.8 |
| Model composition |  |  |  |
| Peptide chains | 19 | 24 | 40 |
| Total atoms | 25592 | 50192 | 75784 |
| Protein residues | 1944 | 2970 | 4914 |
| RNA residues | 472 | 1232 | 1704 |
| Ligands: Mg<sup>2+</sup> / Zn<sup>2+</sup> | 57 / 2 | 109 / 1 | 166 / 3 |
| Average B-factors (min/max/mean) |  |  |  |
| Protein | 12/80/36 | 4/50/15 | 15/112/46 |
| RNA | 7/117/28 | 3/89/25 | 12/175/52 |
| Ligand | 9/91/19 | 4/38/9 | 13/100/31 |
| Model validation |  |  |  |
| RMSD bonds (Å) | 0.003 | 0.002 | 0.002 |
| RMSD angles (°) | 0.535 | 0.512 | 0.500 |
| All-atom clashscore | 3.96 | 4.02 | 4.34 |
| EMRinger score | 3.29 | 4.30 | 3.50 |
| MolProbity score | 1.49 | 1.36 | 1.39 |
| Ramachandran statistics |  |  |  |
| Favored (%) | 96.97 | 97.81 | 97.69 |
| Allowed (%) | 3.02 | 2.19 | 2.31 |
| Outliers (%) | 0 | 0 | 0 |
| Rotamer outliers (%) | 1.49 | 1.48 | 1.39 |
| RNA validation |  |  |  |
| Sugar pucker outliers (%): | 0 | 0 | 0 |
| Angle/bond outliers (%): | 0 | 0 | 0 |
| Bond outliers (%): | 0 | 0 | 0 |

## Notes

### Competing Interest Statement

The authors have declared no competing interest.

